# Alertness fluctuations during task performance modulate cortical evoked responses to transcranial magnetic stimulation

**DOI:** 10.1101/155754

**Authors:** Valdas Noreika, Marc R. Kamke, Andrés Canales-Johnson, Srivas Chennu, Tristan A. Bekinschtein, Jason B. Mattingley

**Affiliations:** Queensland Brain Institute, University of Queensland, St Lucia, QLD 4072, Australia; Department of Psychology, University of Cambridge, Cambridge, CB2 3EB, United Kingdom; School of Computing, University of Kent, Medway, United Kingdom; Department of Clinical Neurosciences, University of Cambridge, Cambridge, United Kingdom; School of Psychology, University of Queensland, St Lucia, QLD 4072, Australia; Canadian Institute for Advanced Research (CIFAR)

**Keywords:** alertness, cortical reactivity, inter-trial variability, motor evoked potentials, transcranial magnetic stimulation

## Abstract

Transcranial magnetic stimulation (TMS) has been widely used in human cognitive neuroscience to examine the causal role of distinct cortical areas in perceptual, cognitive and motor functions. However, it is widely acknowledged that the effects of focal cortical stimulation on behaviour can vary substantially between participants and even from trial to trial within individuals. Here we asked whether spontaneous fluctuations in alertness can account for the variability in behavioural and neurophysiological responses to TMS. We combined single-pulse TMS with neural recording via electroencephalography (EEG) to quantify changes in motor and cortical reactivity with fluctuating levels of alertness defined objectively on the basis of ongoing brain activity. We observed rapid, non-linear changes in TMS-evoked neural responses – specifically, motor evoked potentials and TMS-evoked cortical potentials – as EEG activity indicated decreasing levels of alertness, even while participants remained awake and responsive in the behavioural task.

**IMPACT STATEMENT:** A substantial proportion of inter-trial variability in neurophysiological responses to TMS is due to spontaneous fluctuations in alertness, which should be controlled for during experimental and clinical applications of TMS.

## INTRODUCTION

Transcranial magnetic stimulation (TMS) is a widely used tool for probing human brain function, with applications ranging from the characterization of inter-hemispheric motor cortex interactions and establishment of causal links between neural oscillations and attention, to the clinical treatment of depression and other diseases (Dugué & VanRullen, 2017; Valero-Cabré et al., 2017; Ziemann, 2017). A number of neurophysiological indices of cortical TMS perturbation have been used to contrast experimental conditions of interest, including motor evoked potentials (MEPs) recorded from peripheral muscles (Barker et al., 1985; Bestmann & Krakauer, 2015), and TMS-evoked potentials (TEPs) which are thought to reflect the reactivity of underlying cortical circuits (Chung et al., 2015; Ilmoniemi et al., 1997). These and other outcome measures show varying sensitivity to different experimental manipulations, as well as confounding factors. Perhaps the largest within-participant variation in motor and cortical responses to TMS are observed when contrasting wakefulness and sleep. As healthy adult participants fall into slow wave sleep, MEP amplitude diminishes (Bergmann et al., 2012; Hess et al., 1987; Grosse et al., 2002), whereas TEP amplitude increases in association with a breakdown of effective connectivity, reflecting a state shift from a global to a more local or stereotypical mode of processing (Massimini et al., 2005). Likewise, sleep pressure has been shown to modulate TMS responses during normal waking in daytime hours. For example, MEP-derived motor thresholds are elevated following sleep deprivation (De Gennaro et al., 2007), whereas TEP amplitude increases throughout the day as a function of a natural build-up of sleep pressure (Huber et al., 2013). To date, however, it is not known whether the effects of TMS on neural activity are influenced by fluctuations in the level of alertness that occur during wakefulness. Here we combined single-pulse TMS with concurrent EEG recording and a simple behavioural task to quantify changes in motor and cortical reactivity with fluctuating levels of alertness defined objectively on the basis of ongoing brain activity.

In most studies that have employed TMS to perturb or modify brain activity, behavioural or physiological data are typically collected over experimental sessions that last up to an hour or more (e.g. Darmani et al., 2018; Herring et al., 2015; Salminen-Vaparanta et al., 2013). The implicit assumption is that participants’ levels or alertness remain relatively consistent for the duration of the testing session. This assumption has recently been challenged by findings from brain imaging studies. For instance, Tagliazucchi and Laufs (2014) found that 30% of research participants drifted into a drowsy state (N1 sleep) during resting-state functional magnetic imaging (fMRI) protocols after only three minutes. These periods of early N1 sleep during passive resting-state scans were accompanied by increased signal variance in sensory and motor cortices, increased temporal-temporal and temporal-frontal connectivity, and decreased thalamic-frontal connectivity patterns (Tagliazucchi & Laufs, 2014), highlighting a pervasive source of variance in neuroimaging data due to drowsiness. Likewise, using an active decision-making task, De Gee et al. (2017) demonstrated that brainstem-controlled inter-trial fluctuations in phasic arousal are accompanied by modulation in the involvement of prefrontal and parietal cortices in choice encoding, again implying that fluctuating levels of alertness are potentially a significant source of variability in neural activity.

Further evidence for the contribution of fluctuating levels of alertness to variability in neural activity has come from studies of MEP amplitudes, which tend to be highly variable from trial to trial even when the intensity of the TMS pulses is held constant (Ellaway et al., 1998; Maeda et al., 2002). Several studies have shown that a significant portion of this variance is related to EEG oscillatory activity in a pre-TMS time window (Mäki & Ilmoniemi, 2010; Sauseng et al., 2009; Zarkowski et al., 2006). In particular, trials with higher pre-stimulation alpha power tend to be associated with lower MEP amplitude (Sauseng et al., 2009; Zarkowski et al., 2006). To the extent that increases in alpha power are associated with decreased levels of alertness in participants who are awake and have their eyes open (Gharagozlou et al., 2015; Kaida et al., 2006), spontaneous fluctuations in alertness could be a significant source of inter-trial MEP variability. Unfortunately, previous TMS investigations have not measured or controlled for changes in alertness in their participants, and so it remains unknown whether fluctuations in alertness are systematically associated with changes in TMS-evoked neural activity.

In the present study, we characterize effects of spontaneous fluctuations in alertness on neurophysiological indices of TMS perturbation over the primary motor cortex during a single daytime session. We had four goals: (1) to estimate the latency and stability of fluctuations in alertness over the course of an active, single-pulse TMS session; (2) to test whether fluctuations in alertness modulate the occurrence and amplitude MEPs; (3) to determine whether the amplitude of TEP responses within the first 50 ms after a TMS pulse changes across different levels of alertness; and (4) to assess whether inter-trial variance of MEP and TEP amplitudes is altered with decreases in alertness.

## RESULTS

### Study outline

Participants (N=20) relaxed with eyes closed in a reclining chair, and received single pulses of TMS over the right motor cortex (targeting the first dorsal interosseous (FDI) muscle of their left hand) at 9 different intensities centred on their individual motor threshold (see Materials and Methods for a detailed outline of the experimental protocol). Participants were asked to press a mouse button with their right hand (which was not targeted) after each TMS pulse if they felt a tactile sensation or twitch in their targeted left hand (Fig. 1A-B). At the commencement of the study, participants were informed that they were free to fall asleep but that they should otherwise continue performing the task. They were gently awakened if they failed to respond after 3 successive TMS pulses.

**Figure 1.**
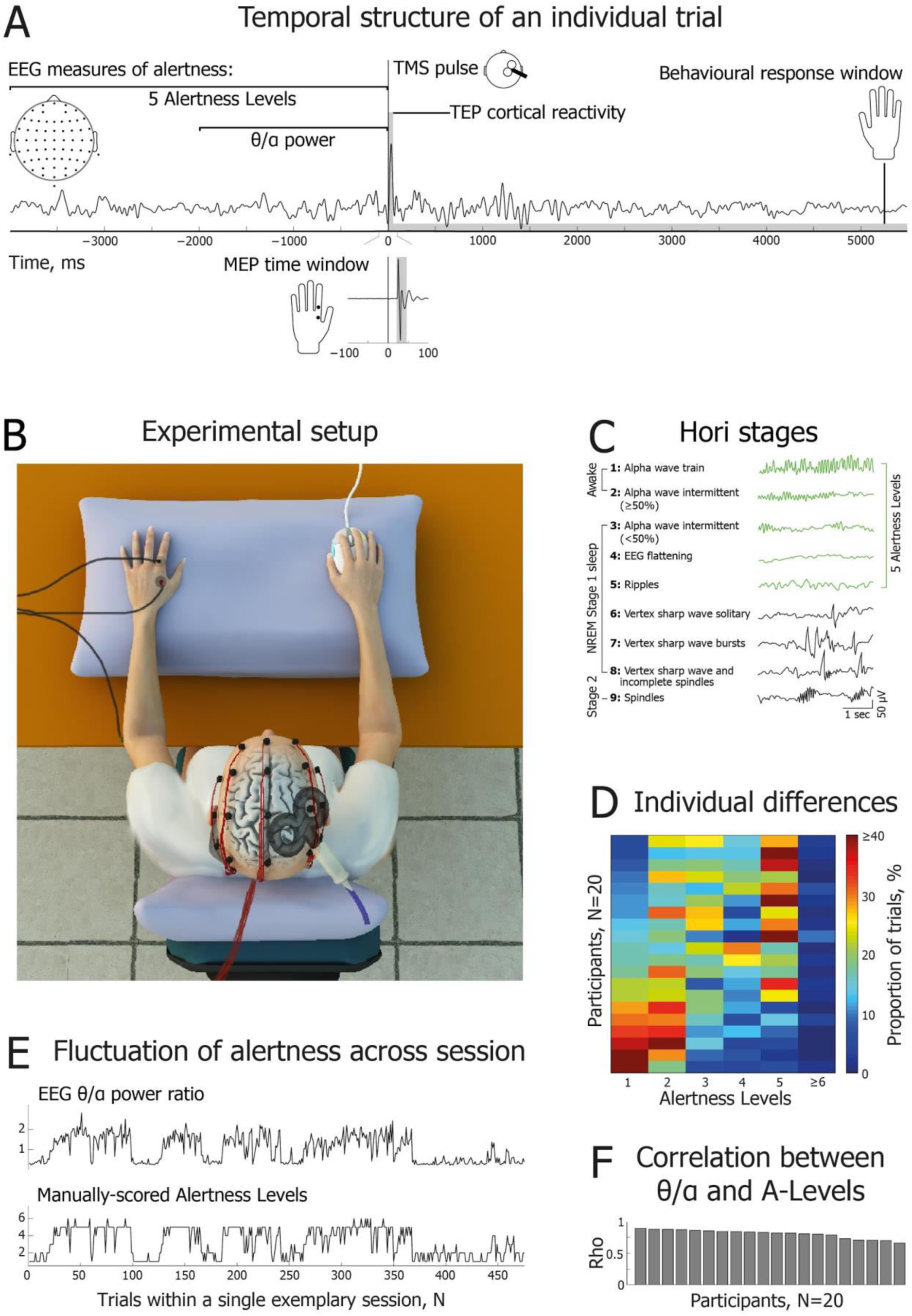
Experimental design and measurements of alertness. (A) Temporal structure of an individual trial. Two EEG windows preceding single-pulse TMS were used to assess alertness: a 4 s window was used for manual scoring of Hori stages (5 Alertness Levels), and a 2 s window was used for automatic calculation of the theta (3-7 Hz) to alpha (8-12 Hz) spectral power ratio. Following each TMS pulse delivered over the right motor cortex, motor evoked potentials (MEPs) were recorded from the first dorsal interosseous (FDI) muscle of the left hand, and TMS-evoked potentials (TEPs) were recorded using high density EEG to characterize cortical reactivity. (B) Schematic of experimental set-up, showing EMG, EEG, TMS and response mouse in situ. (C) Brief definitions and EEG examples of 9 Hori stages of sleep onset, progressing from relaxed wakefulness (Hori Stage 1) to NREM Stage 2 sleep (Hori Stage 9) (modified with permission from Ogilvie (2001)). In the current study, Alertness Levels 1-5 (marked in green) correspond to Hori Stages 1-5. (D) Percentage of trials obtained within each Alertness Level, shown separately for the 20 participants. Datasets are sorted from the most alert participants (lower rows) to the drowsiest participants (upper rows). There were very few epochs of Alertness Level 6 or above. (E) Representative dataset for one participant, showing good agreement between the two EEG measures of alertness across the whole testing session. The upper subplot indicates θ/α ratio; the lower subplot shows fluctuations of Alertness Levels on the Hori scale. (F) Cross-validation of EEG measures of alertness: intra-individual correlations between θ/α power and Alertness Levels across single session trials. Bars represent intra-individual Spearman’s rank order correlation coefficients for the 20 participants, sorted from the most to the least positive coefficients.

To assess the instantaneous level of alertness, a two-fold EEG analysis was applied over the time window immediately preceding each TMS pulse: (1) a binary definition of awake and drowsy states following EEG spectral power signatures (θ/α) averaged across all EEG electrodes (Bareham et al., 2014), and (2) a dynamical definition of alertness levels following a detailed sub-staging system for scoring the transition to N1 sleep (Hori et al., 1994) (Fig. 1C). Both measures were highly correlated (see Fig 1F).

### Fluctuation of alertness during single TMS sessions

On average, TMS sessions lasted for 92.5 min (SD=7, Range=73.5-104.3 min) including time spent switching TMS coils and allowing breaks for participants. During this period, all 20 participants reached Alertness Level 5 or higher, reflecting deep drowsiness with a dominance of EEG theta ripples (see Fig 1C). Notably, it took only 9.44 min on average for participants to reach Alertness Level 5 (SD=8.95, Range=2.25-33.35 min), indicating a rapid decrease of alertness despite the fact that participants were receiving TMS pulses and generating task-specific motor responses.

All participants ceased responding at some point during the testing session, after which they either woke up spontaneously due to TMS, or they were awoken by an experimenter after 3 consecutive unresponsive trials. On average, 13.1% of trials were categorised as “unresponsive” (SD=9.7, Range=2.83-40.6), suggesting a notable impact of drowsiness on task performance. Alertness Levels and unresponsive trials tended to be spread across the testing session, i.e., participants tended to “oscillate” between awake and drowsy states (see Fig 1E and S1). Consequently, there was no systematic increase or decrease in Alertness Level within a session at the group level. Four participants showed a significant but weak positive correlation between Alertness Level and trial number (mean rho=0.21), 6 participants showed a significant negative association (mean rho=-0.27), while the remaining 10 participants showed no significant correlation between Alertness Level and trial number (mean rho=-0.008) (see Fig S1). These results suggest that a given participant’s level of alertness cannot be assumed to decrease continuously over a testing session. Only concurrent EEG measures of alertness can definitively determine a participant’s moment-to-moment level of alertness.

### Fluctuating levels of alertness modulate MEPs

We first assessed corticospinal excitability as a function of alertness and TMS intensity. To this end we calculated the proportion of trials with MEP peak-to-peak amplitude above 50 μV for each of the 9 TMS intensities, separately for the θ/α-defined awake and drowsy trials. A sigmoid function was then fitted across alertness conditions for each participant. The slope of the MEP sigmoid was slightly but significantly shallower in drowsy compared with awake trials (Wilcoxon signed-rank test: z-score=2.02, p=0.044, r=0.32), suggesting mildly increased noise and instability in corticospinal processing (see Fig. 2A; individual participant results are shown in Fig. S2). By contrast, the MEP sigmoid threshold did not differ between awake and drowsy trials (t(19)=1.31, p=0.21, d=0.13, Bayes Factor in favour of the null=2.04).

**Figure 2.**
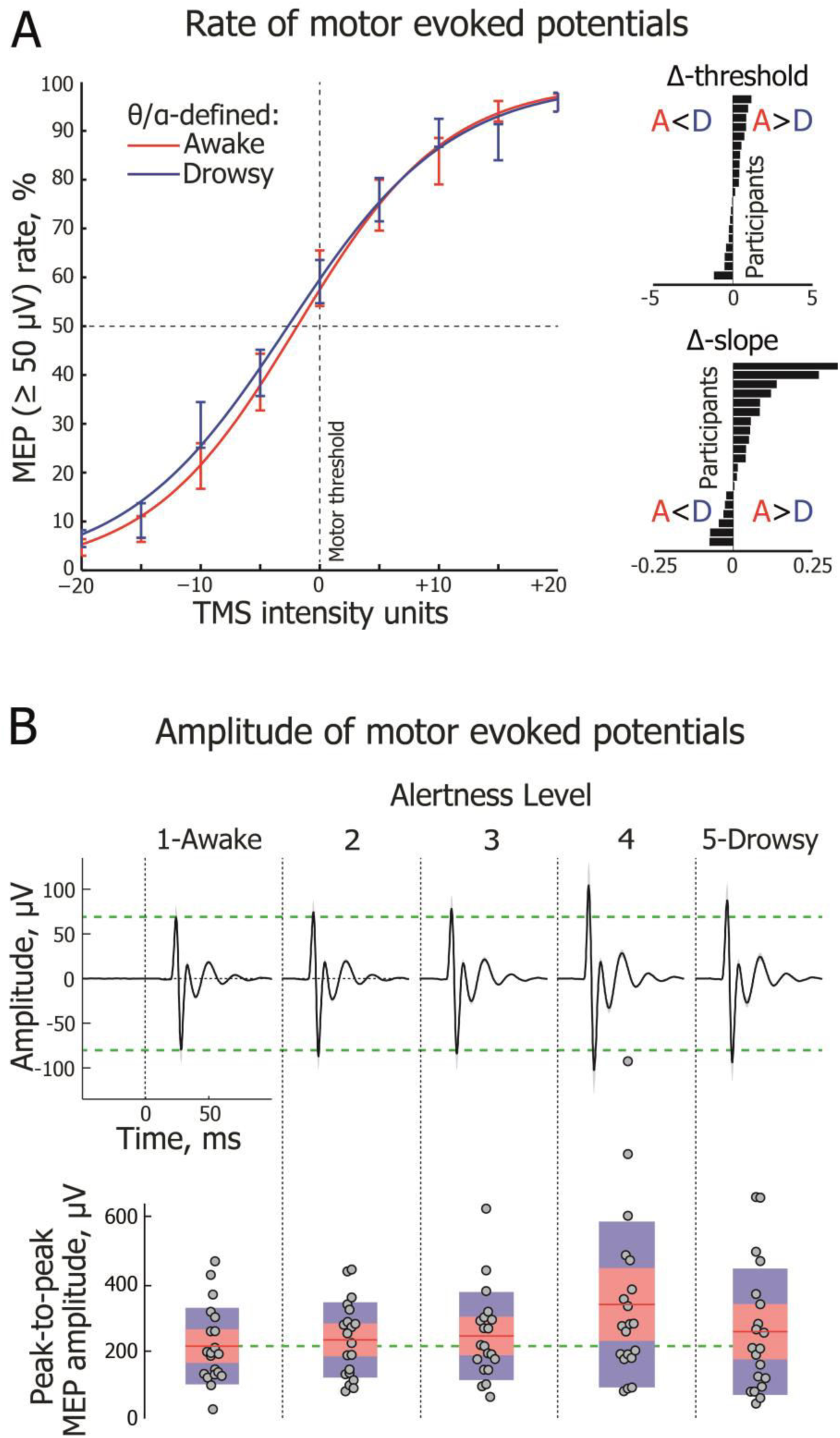
Motor evoked potentials (MEPs) shown across different levels of alertness. (A) Group-averaged frequency of trials with MEPs above a threshold value of 50μV across 9 TMS intensities, centred on individual motor thresholds (0%). Sigmoidal functions are fitted separately to the θ/α-defined awake (red) and drowsy (blue) conditions (error bars represent one standard error of mean, SEM). Insets on the right depict each participant’s sigmoid threshold and slope difference (Awake minus Drowsy), with horizontal bars sorted in ascending order. Only responsive trials are included in the analysis shown in this and other subplots. Alertness states are distinguished here using the EEG θ/α measure taken from a 2000 ms window immediately prior to the TMS pulse. (B, upper panel) Group-level dynamics of MEPs (averaged across three TMS intensities centered on motor threshold) across Alertness Levels 1-5. Horizontal green dashed lines delineate peaks at 25 and 29 ms post-TMS (0 ms) for Alertness Level 1. (B, lower panel) Change in MEP peak-to-peak amplitude across Alertness Levels 1-5. Circles represent individual participants. For each Alertness Level, the red line depicts the group-level mean of peak-to-peak amplitude. The pink shaded region represents 1 standard deviation (SD), and the blue shaded region represents the 95% confidence interval of the mean.

We considered whether the observed difference in MEP slopes was specifically related to alertness, as the amplitude of pre-stimulus alpha oscillations has also been implicated in fluctuations in attention and sensory gating. In these cases, EEG alpha effects are typically evident only within a relatively short pre-stimulus time period of a few hundred milliseconds (Romei et al., 2008), and are restricted to sensory or fronto-parietal regions (Capotosto et al., 2009; van Dijk et al., 2008). Contrary to this, the difference observed here in MEP sigmoid slope as a function of EEG θ/α power was temporally and spatially widespread (Fig. S3).

We next compared MEP peak-to-peak amplitudes between Alertness Level 1, reflecting relaxed wakefulness, and Alertness Levels 2-5, reflecting increasing levels of drowsiness. As shown in Figure 2B, there was a significant increase in MEP amplitude between Alertness Levels 1 and 4 (t(19)=3.5, p=0.0096, d=0.64), as well as an intermediate stepped increase across Alertness Levels 2 (d=0.15) and 3 (d=0.23), and a small decrease at Alertness Level 5 (d=0.27). A linear trend of increasing MEP amplitude was observed across Alertness Levels 1-4 (F(1,19)=11.55, p=0.003, partial η2=0.38), but this was no longer significant when Level 5 was also included (F(1,19)=2.11, p=0.165, partial η2=0.1). These findings indicate a reliably non-linear reorganization of corticospinal excitability at a time when drowsy participants are still conscious and responsive. The most noticeable change in dynamics occurred with the disappearance of alpha waves, at a point where there was EEG flattening and the first occurrence of EEG theta-range ripples, i.e., Alertness Levels 4 and 5 – despite the fact that participants were still responding behaviourally in the task. These observations suggest a much earlier modulation of corticospinal excitability in the initial moments of drowsiness than has been reported previously in studies of MEP changes with sleep deprivation or during NREM sleep (De Gennaro et al., 2007; Grosse et al., 2002; Manganotti et al., 2004).

### Fluctuating alertness modulates TMS-evoked potential (TEP) amplitude

We next assessed post-TMS cortical reactivity measured as TMS-evoked potentials (TEPs) within the first 40 ms after each pulse. Early TEP amplitude is known to increase in response to homeostatic sleep pressure (Huber et al., 2013) and during NREM sleep (Massimini et al., 2005), likely reflecting a combination of synaptic strengthening, changes in neuromodulation, and impaired inhibition (Huber et al., 2013). We hypothesized that, as with MEP amplitude, TEPs should be affected by the level of alertness in drowsy but responsive participants.

Comparing TEP amplitudes between θ/α-defined awake and drowsy trials revealed a reliable increase in cortical reactivity in drowsy trials in a time window from 26-36 ms after the TMS pulse (t(19)=4.02, p=0.00074, d=0.49) (see Fig 3A). This pattern was evident in 18/20 participants (see Fig. S4). While displaying a wide fronto-central spread, the peak difference between awake and drowsy states occurred over the right motor region, directly beneath the TMS coil (see Fig 3B-C). Additional control analyses confirmed that the observed TEP increase for drowsy trials was not due to TMS-evoked scalp muscle artefacts (see Fig S5). We further compared TEP amplitudes at the site of stimulation across Alertness Levels 1-5. As hypothesized, TEP amplitude increased as participants became more drowsy (Mann-Kendall trend test: z=4.7, p=0.0000025, tau=0.77) (Fig 3D-E). Planned comparisons revealed a significant increase in TEP amplitude between Alertness Levels 1 and Level 3 (t(19)=4.54, p=0.00088, d=0.5), 1 and 4 (t(19)=4.38, p=0.00099, d=0.68), and 1 and 5 (t(19)=3.43, p=0.0056, d=0.6). These findings provide the first direct evidence for an inverse association between cortical reactivity and alertness, suggesting that sleep-related changes in neural activity may intrude early in the transition between wakefulness and sleep, while participants are still able to respond in an ongoing behavioural task. Strikingly, the TEP effects emerged at a relatively early Alertness Level, before the appearance of drowsiness ripples or early slow waves (Hori et al., 1994).

**Figure 3.**
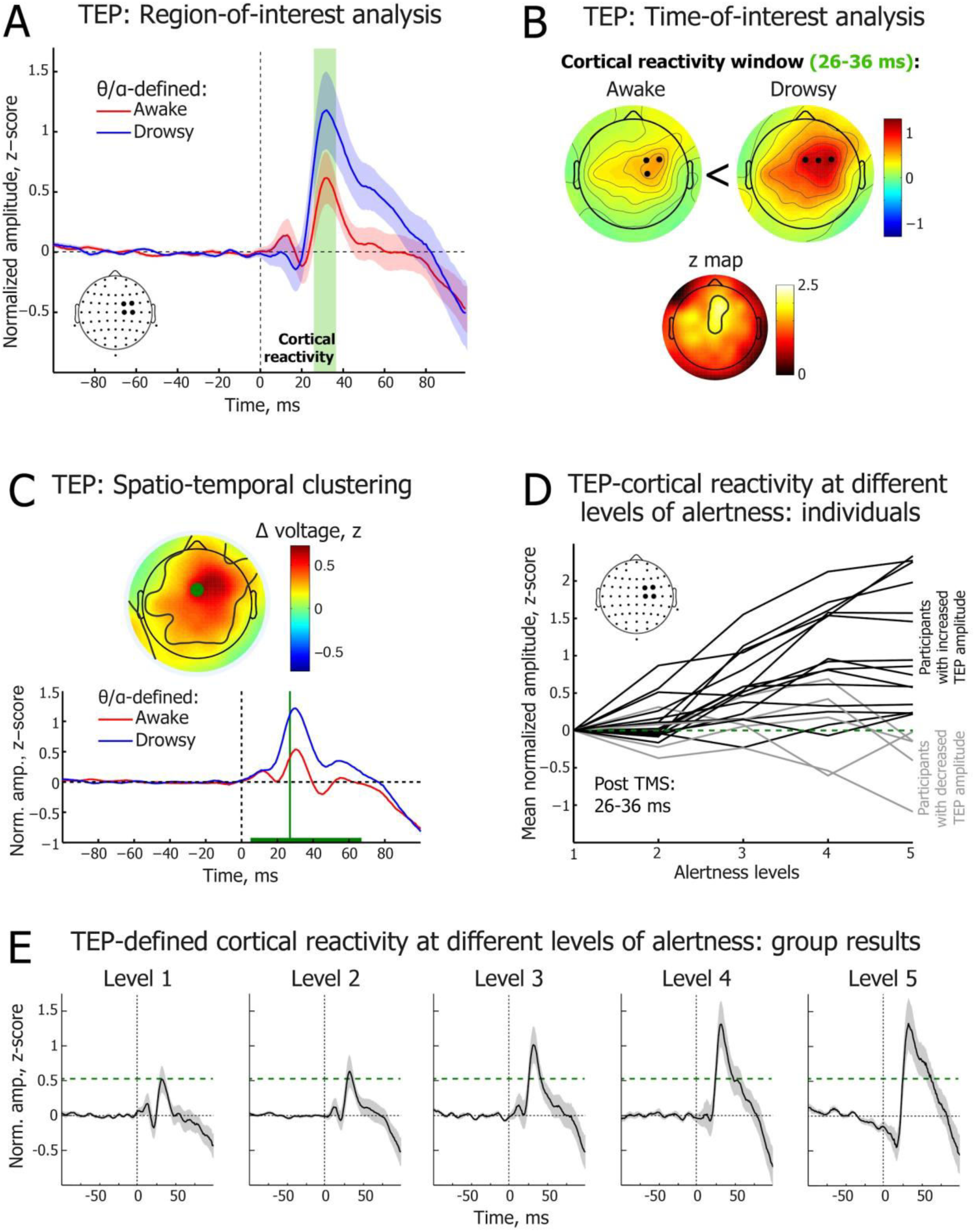
Transcranial magnetic stimulation-triggered cortical reactivity potentials (TEPs) across different levels of alertness. (A) Time course of electroencephalography (EEG) potentials averaged over 4 EEG electrodes beneath the TMS coil in the θ/α-defined awake (red) and drowsy (blue) trials. Green shaded area highlights the cortical reactivity time window (26-36 ms). Only behaviourally responsive trials are included in the analysis shown in this and other subplots. 0 ms corresponds to the time of the TMS pulse. Red and blue shading depicts standard error of the mean (SEM). (B) Topographical distribution of the early TEP mean peak at 26-36 ms post-TMS pulse in the θ/α-defined awake (upper left) and drowsy (upper right) states. Black dots indicate locations of three EEG electrodes with the maximal amplitude in the map. Non-parametric z map (below) reveals region reliably different between awake and drowsy states. (C) 0-100 ms data-driven spatiotemporal clustering of EEG potentials post-TMS pulse between θ/α-defined awake (red) and drowsy (blue) states. TEP amplitude was significantly higher in drowsy trials in a 5-67 ms time window (cluster peak: 27 ms, t = 4884.47, p = 0.004). The green horizontal line depicts the time window of significant difference. The electrode with the largest difference between awake and drowsy states is marked as a green dot in the topographic voltage map, and its waveforms are plotted below. The black contours within the map show the electrodes with statistically reliable differences (cluster). The topographic voltage map is at the peak difference between awake and drowsy states. (D) Individual-level dynamics of TEP cortical reactivity peak amplitude across Alertness Levels 1-5 (TEP amplitude averaged over 26-36 ms across 4 electrodes beneath the TMS coil). Normalized amplitude is shown relative to Alertness Level 1 (green dashed line). Black lines represent participants with higher TEP amplitude at Alertness Level 5 relative to Alertness Level 1 (N=15); grey lines represent participants with lower TEP amplitude at Alertness Level 5 relative to Alertness Level 1 (N=5). (E) Group-level dynamics of TEP waveforms across Alertness Levels 1-5 (TEPs averaged over 4 electrodes beneath the TMS coil). Horizontal green dashed line delineates TEP cortical reactivity peak at 31 ms post-TMS at Alertness Level 1.

### Single-trial MEP and TEP variability across different levels of alertness

Having examined participant-level changes in MEP and TEP amplitudes with spontaneous fluctuations in alertness, we next carried out single-trial analyses of TMS-evoked response variability by pooling data across participants, separately for each Alertness Level. Response variability was quantified as the absolute difference between the TMS-evoked response amplitude in each trial and the mean amplitude in a given Alertness condition.

The variability of single-trial MEP amplitude changed significantly between Alertness Levels 1-5 (Kruskal-Wallis H(4)=297.1, p=4.47E-63; see Fig 4A). Relative to Alertness Level 1 (Mdn=0.3), MEP amplitude variability decreased at Level 2 (Mdn=0.23, Mann-Whitney Z=11.78, p=4.79E-32, r=0.27), Level 3 (Mdn=0.25, Z=9.65, p=5.02E-22, r=0.22), Level 4 (Mdn=0.25, Z=9.96, p=2.19E-23, r=0.23), and Level 5 (Mdn=0.29, Z=6.54, p=6.23E-11, r=0.15). Interestingly, differences in effect sizes suggest a non-linear, U-shaped relationship between MEP amplitude variability and increasing drowsiness.

**Figure 4.**
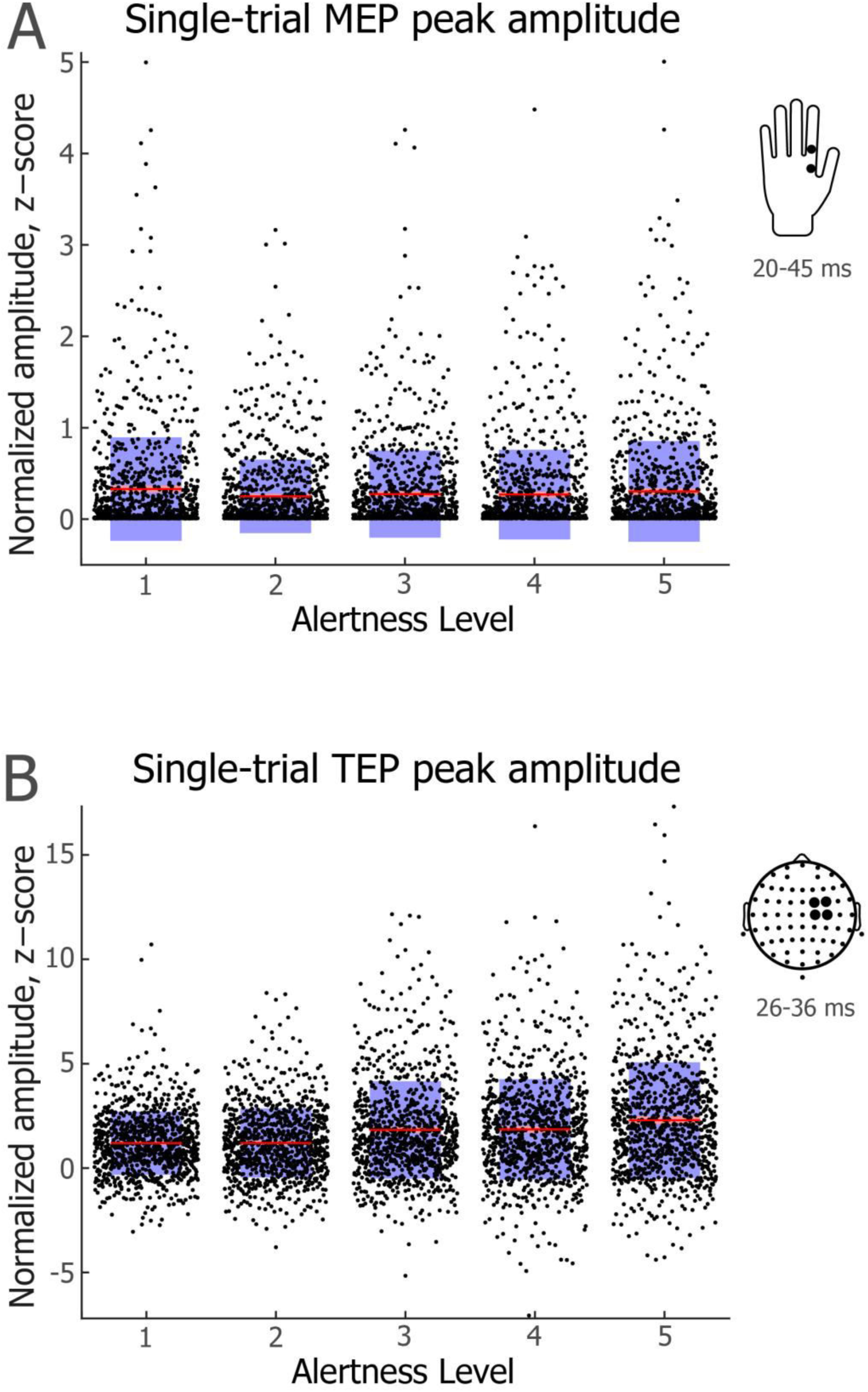
Single-trial MEP and TEP amplitudes across different levels of alertness. (A) MEP peak-to-peak amplitude. (B) TEP peak amplitude. Jittered dots represent individual trials across participants (N=936 per condition). For each Alertness Level, the red line depicts mean amplitude. Pink shading represents 1 standard deviation (SD), which was very small and is thus difficult to discern in the figure. Blue shading represents the 95% confidence intervals for the mean. Insets on the right indicate locations of hand and scalp electrodes and time windows used to detect peak amplitude values.

The variability of single-trial TEP amplitude also changed significantly across Alertness Levels 1-5 (Kruskal-Wallis H(4)=221.3, p=9.9741E-47; see Fig 4B). Relative to Alertness Level 1 (Mdn=0.85), TEP amplitude variability was higher at Level 2 (Mdn=1.03, Mann-Whitney Z=3.12, p=0.0018, r=0.07), Level 3 (Mdn=1.29, Z=8.47, p=2.49E-17, r=0.2), Level 4 (Mdn=1.37, Z=9.19, p=3.96E-20, r=0.21) and Level 5 (Mdn=1.63, Z=12.96, p=2.3E-38, r=0.3). As is evident in Figure 4B, there was a consistent increase in TEP amplitude variability from Alertness Level 1 to Level 5 (standardized Jonckheere-Terpstra trend test statistic=14.65, p=0.00000), suggesting increases in cortical reactivity with decreases in alertness.

### Relationship between single-trial MEP and TEP amplitudes across levels of alertness

In a final step, we asked whether MEP and TEP amplitudes were correlated at a single trial level, and whether any such association was modulated with changes in alertness. There was no significant correlation between MEP and TEP amplitudes at Alertness Level 1 (Spearman’s rho=0.07, p=0.026; not significant after correction) or Level 2 (Spearman’s rho=0.07, p=0.025; not significant after correction). As shown in Figure S6, however, there was a significant positive correlation between MEP and TEP amplitudes at the deeper levels of drowsiness, including Alertness Level 3 (Spearman’s rho=0.22, p=1.73e-11), Alertness Level 4 (Spearman’s rho=0.19, p=3.14e-09), and Alertness Level 5 (Spearman’s rho=0.29, p=9.73e-20) (see Fig. S6). Taken together, these findings indicate that while MEP and TEP responses are independent during wakefulness, they become positively associated with increases in drowsiness, suggesting a gradual shift in state toward a non-differentiated mode of neural processing.

## DISCUSSION

Most studies that use TMS to investigate perceptual, cognitive and motor function in human participants do not consider the possibility that fluctuating levels of alertness across a typical testing session might lead to measurable changes in both behavioural and associated patterns of brain activity. Here we used single-pulse TMS delivered over the right motor cortex while simultaneously measuring MEPs and TEPs across different, objectively defined levels of alertness while participants engaged in a simple tactile perception task. Participants exhibited fluctuating levels of alertness across the testing session, as indexed by continuous EEG recordings, but continued to respond behaviourally even in relatively deep states of drowsiness. Strikingly, both motor evoked responses and TMS-evoked cortical reactivity were altered across different levels of alertness. Specifically, we found that MEP amplitudes peaked during EEG flattening (Alertness Level 4), whereas TEP cortical reactivity increased earlier and remained stable across Alertness Levels 4 and 5. These findings highlight that a substantial proportion of inter-trial variability in neurophysiological responses to TMS can potentially be attributed to spontaneous fluctuation in alertness.

Inter-trial and inter-subject variability in MEP amplitude is a well-known source of data variance in TMS experiments (Kiers et al., 1993; Ellaway et al., 1998; Rösler et al., 2008; Schutter et al., 2010; Sommer et al., 2002), and it has been suggested that 30 or more trials are required to provide a reliable estimate of MEP amplitude (Goldsworthy et al., 2016). The non-stationarity of MEP amplitudes has been attributed to a number of factors, including pre-stimulus voluntary muscle contraction (Kiers et al., 1993), variation in the number of recruited alpha-motor neurons (Rösler et al., 2008), variation in the synchronization of motor neuron discharges (Rösler et al., 2008) and functional hemispheric asymmetries (Schutter et al., 2010). Furthermore, a recent series of studies found that the amplitude or phase of pre-TMS EEG oscillations can predict MEP amplitude, including alpha (Iscan et al., 2016; Sauseng et al., 2009; Schulz et al., 2014; Zarkowski et al., 2006), beta (Mäki & Ilmoniemi, 2010; Keil et al., 2014; Schulz et al., 2014) and gamma (Zarkowski et al., 2006) frequency bands. Extending these studies, our findings suggest that changing levels of alertness could be a key factor in brain state-modulation of corticospinal excitability. For instance, Zarkowski et al. (2006) demonstrated that MEP amplitude is negatively correlated with pre-TMS alpha power (10-13 Hz) and positively correlated with pre-TMS gamma power (30-60 Hz), with an alpha/gamma ratio being the strongest predictor of MEP amplitude. While a theta/alpha ratio can index the level of alertness in eyes-closed experiments, such as in the present study, the alertness-indexing frequencies are shifted upward in eyes-open paradigms (Eoh et al., 2005; Kaida et al., 2006; Zhao et al., 2012). It is therefore likely that the results of Zarkowski et al. (2006) and other studies were influenced by changing levels of alertness in the participant sample.

Our finding of non-linear changes in MEP amplitude with decreasing levels of alertness might explain previous contradictory findings regarding sleep deprivation effects on MEP amplitude. While several studies have reported an increase in corticospinal motor threshold following sleep deprivation (Manganotti et al., 2001; De Gennaro et al., 2007), other studies have failed to find any such effect (Civardi et al., 2001; Manganotti et al., 2006). Arguably, due to individual differences in instantaneous drowsiness levels and potentially different time of day and duration of testing, the dominant level of alertness varied between these studies, confounding their comparison. For instance, datasets with a relatively high proportion of trials obtained during Alertness Level 4 would likely indicate higher MEP amplitude compared with other datasets. Unfortunately, a fine-grained measurement of alertness is seldom undertaken in MEP studies, even when EEG is recorded, e.g., “sleepiness” or NREM Stage 1 sleep are usually treated as a uniform state (Manganotti et al., 2004), even though a more detailed analysis can reveal at least 6 micro-states within N1 sleep (Hori et al., 1994; see Fig 1C).

Regarding the modulation of TEPs with sleepiness, Huber et al. (2013) observed an increase of TEP amplitude as a function of prolonged wakefulness as well as following sleep deprivation. Contrary to our results, however, they found no association between TEP amplitude and short-lasting episodes of drowsiness. This discrepancy is most likely attributable to the fact that Huber et al. (2013) relied solely on a behavioural definition of drowsiness (performance in a visuomotor tracking task), whereas we used fine-grained EEG measures of alertness that could be quantified independently of fluctuations in behaviour. In another recent study, TEP amplitude was found to depend on the interaction between sleep pressure and a phase of circadian cycle rather than sleep homeostasis alone (Ly et al., 2016). Furthermore, the same study found that an increase in TEP amplitude was associated with an increase in EEG theta power across 29 h of sustained wakefulness. Unfortunately, the authors did not report whether such an association held over a shorter period of time (e.g., 45 - 90 mins), as would be the case in typical TMS studies.

Sleep-related increases in TEP amplitudes likely reflect facilitation of a stereotypical, local mode of processing, which has been linked to the loss of consciousness (Massimini et al., 2005, 2012; Tononi & Massimini, 2008). Similarly, an increase in TEP amplitude is observed during dreamless xenon anaesthesia, whereas dreamful ketamine anaesthesia shows no changes in TEP amplitude compared with wakefulness, suggesting a specific link between TEP amplitude and (un)consciousness (Sarasso et al., 2015). Contrary to these suggestions, however, here we found that the perturbational TEP marker of (un)consciousness arose at an early stage of drowsiness, when participants were still able to respond behaviourally and well before they fell asleep. Thus, we would suggest that changes in local cortical processing as indexed by TEPs primarily reflect different levels of alertness rather than the presence or absence of sensory awareness.

An increase in stereotypical local processing with increasing drowsiness was also observed in the association between TEPs and MEPs. Fecchio et al. (2017) showed that trials with a high MEP amplitude were also associated with a relatively high TEP amplitude compared with trials with a low MEP amplitude. In the present study, we found a gradually increasing inter-trial variability of TEP amplitude across decreasing levels of alertness. By contrast, variability in MEP amplitude showed a more complex U-shaped pattern characterised by an initial decrease during early stages of drowsiness and a subsequent increase with higher levels of drowsiness, suggesting potentially independent neural pathways in the transition between wakefulness and sleep. Nevertheless, while TEP and MEP amplitudes showed no correlation at Alertness Levels 1 and 2, these two neural indices became positively correlated with increasing drowsiness, i.e. at Alertness Levels 3-5. These findings suggest that within-trial associations between corticospinal excitability (MEP) and cortical reactivity (TEP) depend on the level of alertness, which should be controlled for when experiments involve hundreds of trials delivered over a prolonged testing session (Fecchio et al., 2017; Helfrich et al., 2012).

Several strategies exist for dealing with changing levels of alertness during behavioural testing. If alertness decreases throughout a session, an additional factor of trial or block number could be added to a statistical model as an alertness regressor or covariate. However, we found that an initial decrease in alertness did not persist throughout the testing session, and participants tended to “oscillate” between awake and drowsy periods (see Fig S1), which precludes any straightforward inference of decreasing alertness over the course of a single testing session. Alternatively, reaction times (RTs) could be used as a behavioural index of alertness in active TMS experiments, as RTs typically lengthen with decreases in vigilance (Schmidt et al., 2009), increases in drowsiness (Ogilvie & Wilkinson, 1984), and following sleep deprivation (Ratcliff & Van Dongen, 2011). However, such a strategy would not be possible in passive paradigms in which participants are not required to respond (Gordon et al., 2018; Massimini et al., 2005). Furthermore, as both trial counts and reaction times are relative measures, participants who maintain high alertness throughout a session could be falsely labelled as drowsy in a proportion of trials. Arguably, concurrent EEG recording should be the gold standard for assessment of single-trial alertness, as it provides quantifiable and reliable signatures of instantaneous brain-states, including alpha- and theta-derived Hori stages of sleep onset (Hori et al., 1994). As an alternative to the tedious manual scoring of Hori stages (Alertness Levels), an automated EEG method based on wakefulness and sleep grapho-elements is available for the detection of drowsiness from EEG data (Jagannathan et al., 2018). While methods that weight the dominance of EEG theta and alpha oscillations are suitable for eyes-closed paradigms (Hori et al., 1994; Jagannathan et al., 2018), such as resting state or phosphene studies (Bonnard et al., 2016; De Graaf et al., 2017), the power of higher EEG frequencies should be considered when assessing alertness during active eyes-open experiments (Eoh et al., 2005; Kaida et al., 2006; Zhao et al., 2012). Finally, when EEG measurements are not available or feasible, TMS experiments could be carried out in short blocks of just a few minutes each (e.g., 3-5 min) and inter-block intervals could be used to assess instantaneous subjective sleepiness, for example by asking participants to undertake the 9-graded Karolinska Sleepiness Scale (Åkerstedt & Gillberg, 1990; Kaida et al., 2006).

Future studies should compare other methods for capturing the contribution of fluctuating levels of alertness to data variance in TMS studies. In addition, follow up studies are needed to assess the impact of changing levels of alertness on MEP and TEP amplitude in eyes-open paradigms. It will also be important to investigate whether neurophysiological effects of decreasing alertness are limited to individuals who are likely to fall asleep in a situation of prolonged inactivity, such as our participants. If this turned out to be the case, it might be advisable to selectively recruit individuals who are very unlikely to fall asleep during daytime hours to TMS experiments that require a high and stable level of alertness.

To conclude, our findings challenge the widely held assumption that the cortex is maintained in a more or less “steady state” when participants undertake experimental investigations of perceptual, cognitive or motor function. Our findings demonstrate that spontaneously occurring fluctuations in alertness differentially modulate cortical reactivity over relatively short durations, even when participants are tested during day time hours when they would normally be awake and performing typical activities of daily living. Our study highlights the importance of controlling for spontaneous fluctuations in alertness at a single trial level in non-invasive brain stimulation studies.

## MATERIALS AND METHODS

### Participants

Twenty participants (7 male; mean age 23.7 years: age range 21-33 years) took part in the study. All participants were screened for contraindications to TMS (Rossi et al., 2009), which included having no history of hearing impairment or injury, and no neurological or psychiatric disorders. All participants were right handed, as assessed using the Edinburgh Handedness Scale (Oldfield, 1971). The mean handedness index was 0.79 (SD=0.19; range 0.3 to 1). Potential participants were also screened with the Epworth Sleepiness Scale (ESS) (Johns, 1991). The mean ESS score was 9.4 (SD=4.3), which indicates that most of the participants had a slight to moderate chance of dozing off in a situation of prolonged inactivity.

The experimental protocol was approved by the Medical Research Ethics Committee of The University of Queensland (UQ), and the study was carried out in accordance with the Declaration of Helsinki. All participants gave informed, signed consent. Participants were recruited through an electronic volunteer database managed by UQ’s School of Psychology. They received $30 for taking part in the study. There were no adverse reactions to TMS.

### Electromyography (EMG)

Surface EMG was recorded from the first dorsal interosseous (FDI) of the left and right hands using disposable 24 mm Ag-AgCl electrodes (Kendall H124SG by Covidien; MA, USA). The electrodes were placed in a belly-tendon montage with the reference over the proximal phalanx of the index finger and a common reference on the right elbow. Raw EMG signals were amplified (×1000) and filtered (20-2000 Hz; 50 Hz notch filter) using a Digitimer NeuroLog system (Digitimer; Hertfordshire, UK). The data were digitised at 5000 Hz using a Power 1401 and Signal (v5) software (Cambridge Electronic Design; Cambridge, UK) and stored for offline analysis on a PC. Throughout the experiment EMG activity was monitored on-line using a digital oscilloscope with a high gain. Participants were prompted to relax if any unwanted muscle activity was observed.

### Transcranial magnetic stimulation (TMS)

TMS was applied to the right primary motor cortex using a Magstim 2002 stimulator and a 70 mm figure-of-eight coil (#9925-00; The Magstim Company; Carmarthenshire, UK). The site for stimulation was the point on the scalp over the motor cortex that elicited the largest and most consistent amplitude MEPs from the left FDI. This stimulation ‘hotspot’ was found by placing the TMS coil tangentially on the scalp with the handle pointing posteriorly and laterally at ∼45° to the sagittal plane, and stimulating at an intensity that was assumed to be slightly suprathreshold for most individuals. Once the hotspot had been identified it was marked using an infrared neuro-navigation system (Visor 2 by ANT Neuro; Enschede, The Netherlands). A small piece of foam (∼ 5) mm thick was then placed under the centre of the TMS coil so that it was not in physical contact with any EEG electrodes. The hotspot was re-marked and the location and orientation of the TMS coil were maintained throughout the testing session with the aid of the neuro-navigation system. Accuracy of coil position and handle orientation were kept within 5 mm and 5 degrees, respectively, but were typically within 3 mm and 3 degrees. Resting motor threshold was determined using the relative frequency method with a criterion of ≥50 µV (peak-to-peak) MEP amplitude in at least five out of ten consecutive trials (Ikoma et al., 1996; Rossini et al., 1994; Samii et al., 1996). A two-down, one-up staircase was used, starting at a suprathreshold intensity. Mean motor threshold for the group was 53.1% (Range 34 – 74%) of maximal TMS output intensity.

### Electroencephalography (EEG)

Continuous EEG data were acquired using a 64 channel BrainAmp MR Plus amplifier, TMS BrainCap and Brain Vision Recorder (v1) software (Brain Products; Gilching, Germany). A high chloride abrasive electrolyte gel was used (Abralyt HiCl by Easycap; Herrsching, Germany) and electrode placement corresponded with the International 10-10 system. Data were sampled at 5 kHz with a bandpass filter of DC-1000 Hz and resolution of 0.5 mV (± 16.384 mV). Recordings were referenced online to the left mastoid, and electrode impedance was typically kept below 5 kΩ.

### Experimental procedure

Participants were seated in a comfortable reclining chair that included head and leg support (see Figure 1B). After placing the EMG and EEG electrodes, participants had their eyes blindfolded and the lights in the lab were dimmed. They were instructed to relax for a few minutes while estimation of individual resting motor threshold was performed. Participants’ hands were comfortably supported with pillows. After threshold estimation the combined TMS-EEG experiment was carried out, and participants were reminded to stay relaxed and keep their eyes closed. They were also instructed to pay attention covertly to their left hand and to respond by clicking one of the two keys on a mouse held in their right hand if they felt a tactile sensation, such as a twitch or a touch, in their left hand at the time of each TMS pulse (see Figure 1A). Participants were explicitly instructed that they were permitted to fall asleep should they wish to. If no responses were registered after 3-5 consecutive trials (i.e., failure to press a mouse button within 6 seconds of a TMS pulse), participants were gently awakened verbally and reminded to continue the task.

During stimulation, nine TMS intensities centred on the individual resting motor threshold were used (−20%, −15%, −10%, −5%, 0%, +5%, +10%, +15%, +20%). Given that TMS stimulator output intensity is measured in whole numbers from 1 to 100, the calculated percentage from threshold intensity was rounded. This yielded slightly different sized steps from −20% to +20% for some individuals, and this was taken into consideration when fitting sigmoidal functions at the single participant level. For each individual, 520 trials of single pulse TMS were delivered, with an average inter-pulse interval of 9.5 sec and a uniformly distributed random jitter of ±1000 ms. Thus, the inter-pulse interval lasted between 8.5-10.5 s. We incorporated a relatively long inter-pulse interval to facilitate the natural development of drowsiness, and to allow sufficient time for a return of tonic EMG activity to its baseline level. As our aim was to obtain the maximum number of evoked responses (MEPs and TEPs) around the TMS threshold intensity, the following number of trials was delivered at each TMS intensity: 40 trials (7.7% of a total) at each of the −20%, −15%, −10%, +10%, +15% and +20% intensities; 80 trials (15.4% of a total) at each of the −5% and −5% intensities; 120 trials (23.1% of a total) at 0% (i.e., at the individual resting motor threshold). Trial order was randomized throughout the experiment. TMS pulses were delivered in 8 blocks of 65 trials. One experimenter held the TMS coil and the other monitored ongoing EEG; these individuals switched their places after each block. An extended rest was provided after 4 blocks to allow participants a break from the task, to change the heated TMS coil, and to reduce the impedance of any EEG electrodes if required. Data collection lasted approximately 90 minutes. In an effort to reduce the potential impact of any circadian fluctuation in cortical excitability (Sale et al., 2007), all testing sessions commenced at 1.00 pm (a time at which participants were more likely to feel drowsy after having had their lunch).

### Motor evoked potential (MEP) analyses

Peak-to-peak amplitudes of MEPs evoked by TMS pulses delivered over the right motor cortex were calculated for each trial within a 20-45 ms time window using Signal (v5) software (Cambridge Electronic Design; Cambridge, UK). Trials containing phasic muscle activity in the left FDI channel within 100 ms prior to a TMS pulse being delivered were discarded from the analyses.

We characterized modulations of MEPs as a function of alertness levels by fitting a sigmoid function to the proportion of trials that evoked MEPs (constrained from 0 to 1 on the y axis) across the 9 TMS intensities (−20%, −15%, −10%, −5%, 0%, +5%, +10%, +15%, +20). We then compared threshold and slope measures in awake and drowsy trials separately for each participant. A 50 μV cut-off threshold in peak-to-peak amplitude (Ikoma et al., 1996; Rossini et al., 1994; Samii et al., 1996) was used to define the presence of an MEP. A sigmoid function was fitted to each individual participant’s data:

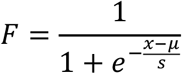

where **F** is the MEP ratio, **x** is the TMS intensity, **µ** is the threshold value (the TMS intensity at the inflection point), and **s** is inversely proportional to the slope at the threshold. The actual slope of the fitted sigmoid was calculated by fitting a straight line between a point 0.1 above the inflection point and a point 0.1 below it (see Figure S2).

To assess dynamics of MEP peak-to-peak amplitude across Alertness Levels 1-5, responsive trials with MEP amplitude at least twice as high as the peak-to-peak distance in the −100-0 ms baseline window were averaged separately for each Alertness Level and each participant. In an effort to control for MEP variance as a function of TMS intensity, only three TMS intensities (−5%, 0%, +5%) around each individual’s motor threshold were included in the analysis of MEP changes across Alertness Levels.

### EEG pre-processing and analysis: pre-TMS spectral power

EEG data pre-processing was carried out using EEGlab toolbox for Matlab (Delorme & Makeig, 2004), with two separate pre-processing pipelines developed for the analysis of EEG activity before and after TMS pulses. To calculate EEG spectral power before TMS, the recordings were downsampled to 500 Hz, and then epoched in −4000 ms to −10 ms time segments preceding each TMS pulse. The noisiest epochs were manually deleted, and the most deviant EEG channels were detected with the ‘spectopo’ function before running the independent component analysis (ICA) for further removal of artefacts such as eye blinks and saccades, heartbeats, and muscle noise. ICA was carried out on relatively clean channels only, whereas the noisy channels were recalculated by spherical spline interpolation of surrounding channels after deleting ICA components with artefacts. Data were again manually inspected and several remaining noisy epochs were deleted. On average, 58 trials (11.2%) were discarded per single participant during EMG and EEG pre-processing, leaving on average 462 trials per participant (SD=38; Range=348-513 trials) for the subsequent analyses.

The spectral power of EEG oscillations over the 4 s time interval immediately preceding each TMS pulse was computed using a Hilbert transform, set from 1.5 Hz to 48.5 Hz in steps of 1 Hz. Given that estimation of spectral power of slow oscillations can be difficult close to the edges of EEG segments (Cohen, 2014), and we were particularly interested in the spectral power just before each TMS pulse, a dummy copy of each EEG epoch was created by flipping the left and the right sides of each pre-TMS epoch along the time axis. The resulting “mirror image” data were the concatenated with the original pre-TMS data; that is, the time axis of the obtained 7.976 s EEG epochs extended from −4000 ms to −12 ms (original) and then back from −8 ms to −4000 ms (mirror). In this manner, an abrupt discontinuity was avoided in the time window just before the TMS pulse, thus enabling a more stable estimate of spectral power. After Hilbert transformation, the “mirror” part of the EEG epoch was deleted, retaining the original pre-TMS window from −4000 ms to −12 ms. To reduce data size, EEG recordings were down-sampled to 250 Hz before running the Hilbert transform.

### EEG measures of alertness

Two complementary EEG measures were used to assess participants’ level of alertness before each TMS pulse: (1) the Hori scoring system of sleep onset EEG (Hori et al., 1994), and (2) a ratio between EEG spectral power of pre-TMS theta and alpha oscillations, which we refer to here as the ‘θ/α’ measure of alertness (Bareham et al., 2014; Noreika et al., 2017).

The Hori system relies on visual scoring of 4 s segments of continuous EEG data (Hori et al., 1994). It consists of 9 stages reflecting a gradual progression from wakefulness to sleep, from Hori Stage 1 which refers to alpha-dominated relaxed wakefulness, to Hori Stage 9 which is defined by the occurrence of complete spindles coinciding with classic Stage 2 NREM sleep (see Fig 1C). The Hori system has been used to map dynamic wake-sleep changes in ERPs (Nittono et al., 1999), EEG spectral power (Tanaka et al., 1997), reaction times, and the rate of subjective reports of being asleep (Hori et al., 1994). In the present study, Hori stages were visually assessed by an experienced sleep researcher (VN) who was blind to participants’ responsiveness and the TMS intensity on any trial. For scoring purposes, only 19 EEG channels of the standard 20-10 system were used (Fp1, Fp2, F7, F3, Fz, F4, F8, C3, Cz, C4, T7, T8, P7, P3, Pz, P4, P8, O1, O2), and EEG recordings were low pass filtered (20 Hz). Previous research has found that participants are typically unresponsive in Hori Stages 6 and above (Ogilvie, 2001), so our analysis was restricted to Hori Stages 1-5, which we refer to here as ***Alertness Levels 1-5***.

Hori Stages 1 to 4 are marked by decreasing activity in the alpha range, and Hori Stages 4 to 8 are characterized by an increase in activity in the theta range (Hori et al., 1994). Thus, progression of drowsiness can be quantified by a ratio of the spectral power of the alpha and theta EEG frequency bands. Specifically, here drowsiness was quantified as a period of time with an increased θ/α ratio of spectral power (Bareham et al., 2014). To apply this measure, theta (4.5-7.5 Hz) and alpha (8.5-11.5 Hz) power was first averaged in time from −2000 ms to −12 ms, and the θ/α ratio was then calculated for each trial and electrode. Next, the θ/α ratio was averaged across all electrodes, resulting in a single “alertness” value per trial. Finally, trials were split into the most strongly “awake” (45%) and most strongly “drowsy” (45%) trials, excluding the 10% of trials that were intermediate between the two extremes.

Importantly, in addition to spontaneous fluctuations in alertness, the spectral power of EEG pre-stimulus oscillations can reflect attentional sampling and/or sensory gating (Capotosto et al., 2009; Romei et al., 2008; van Dijk et al., 2008). We expected that an alertness-related effect would be spatially and temporally widespread and consistent, so we repeated the MEP analysis by splitting the data between awake and drowsy trials separately for each EEG electrode, and in 20 equally sized pre-TMS time bins of 100 ms duration, from −2000 ms to 0 ms relative to the TMS pulse.

### Pre-TMS level of alertness

All participants completed the experimental task and reached the expected Alertness Level 5 or higher, marked by the occurrence and dominance of theta waves. At a group level, a comparable proportion of awake and drowsy trials were obtained as per the criteria defined above (Alertness Levels 1-2: M=45.17%, SD=19.92; Alertness Levels 4-5: M=35.68%, SD=16.44) (see Fig 1D).

Thus, even though the Hori system provides absolute electrophysiological signatures of the depth of drowsiness, the θ/α ratio was used to identify equal proportions of awake and drowsy trials within each participant. Given that the θ/α measure is relative, there was a risk of mislabelling trials for some participants, as it would make a split between “awake” and “drowsy” trials even if all of them happened to be Alertness Level 1. Thus, to verify the use of θ/α data splits, we compared these two measures at an individual level and at the level of the group as a whole. First, we carried out correlation analyses between the two measures of alertness within each participant. Second, we compared correlation coefficients against zero to assess the consistency of association between the Hori and the θ/α measures. At an individual level, Hori and θ/α scores were positively and significantly correlated for all 20 participants (individual rho ranged from 0.66 to 0.9). Group analysis confirmed a very strong association between these two electrophysiological measures of alertness (one sample t test: t(19)=51.99, p<0.000005), confirming that the θ/α ratio was well suited to assessing the level of alertness in the sample here (see Fig 1F).

### EEG pre-processing and analysis: TMS-EEG reactivity

Analysis of EEG reactivity to TMS pulses in the first 50 ms time window requires a perfect alignment of TMS markers with the onset of the actual TMS pulses. Given that there was some delay and jittering between a TMS marker and the pulse itself (M=9.6 ms, SD=1.7 ms), EEG markers indicating TMS intensity were automatically adjusted to the time point of the actual TMS pulse. For this, raw EEG data were segmented ±200 ms around each TMS marker, and global field power (GFP) was calculated as a standard deviation of voltage across all electrodes, resulting in a single time waveform for each TMS marker. Each obtained waveform was baseline corrected to the −200 ms to −50 ms time window, and each time sample was transformed to its absolute value. The remaining time window of −49 ms to +200 ms was scanned, searching for the first time point where a GFP value that exceeded the maximal baseline GFP value by a factor of five, which indicated the onset of a TMS artefact. The TMS marker was then reallocated to this point in the continuous EEG recording.

The EEG data were processed following an ICA-based approach of TMS-EEG artefact cleaning (Rogasch et al., 2014). First, EEG data were segmented from −1000 ms to +1000 ms around the onset of TMS artefact. Next, the segments were baseline corrected to the mean of the interval from −500 ms to −100 ms time window. A line was then fitted to the data from −2 ms to 15 ms, thus deleting the initial TMS-EEG artefact, and the epochs were down-sampled to 1000 Hz. The most deviating EEG channels were then detected with the ‘spectopo’ function and the first round of independent component analysis (ICA) was performed without using noisy channels. After deleting a very distinctive early high amplitude component that reflected TMS-evoked contraction of scalp muscles, EEG data were filtered (1-80 Hz) and epoched from −400 ms to +600 ms around the onset of the TMS marker. Once again, any deviating EEG channels were identified and a second round of ICA was carried out without using noisy channels. Independent components reflecting TMS-EEG decay artefact, eye movements, auditory evoked potentials, 50 Hz line noise, and other sources of noise, were deleted, after which bad channels were recalculated using spherical spline interpolation. The EEG segments were again baseline corrected (−100 ms to −3 ms), and manually inspected to delete epochs that still contained a residual TMS artefact. To account for within-trial variance, raw voltages of each individual trials were transformed to z-scores using the mean and standard deviation of the baseline period (−100 to −3 ms). Trials were then split into different levels of alertness.

To assess changes in EEG reactivity to TMS perturbation as a function of alertness, the four electrodes immediately beneath the TMS coil were chosen to contribute their voltage values to a region of interest (ROI), and these were then averaged within each participant. The group-level waveform was then plotted, revealing an early TEP peak at 31 ms post-TMS pulse. The data were then split between Alertness Levels and the mean amplitude (±5 ms) around the peak (26-36 ms) was calculated for each participant and each level of alertness.

ERP dynamics were additionally studied using data-driven spatiotemporal clustering analyses similar to what we have described previously (Chennu et al., 2013). Awake and drowsy trials were compared in the time windows of interest (15-100 ms) by averaging single-subject data and running group level clustering. Using modified functions of FieldTrip toolbox (Maris and Oostenveld, 2007; Oostenveld et al., 2011), we compared corresponding spatiotemporal points in individual awake and drowsy trials with an independent samples t-test. Although this step was parametric, FieldTrip uses a nonparametric clustering method (Bullmore et al., 1999) to address the multiple comparisons problem. t values of adjacent spatiotemporal points whose p values were less than 0.05 were clustered together by summating their t values, and the largest such cluster was retained. A minimum of two neighbouring electrodes had to pass this threshold to form a cluster, with the neighbourhood defined as other electrodes within a 4 cm radius. This whole procedure, that is, calculation of t values at each spatiotemporal point followed by clustering of adjacent t values, was repeated 1000 times, with recombination and randomized resampling before each repetition. This Monte Carlo method generated a nonparametric estimate of the p value representing the statistical significance of the originally identified cluster. The cluster-level t value was calculated as the sum of the individual t values at the points within the cluster.

We considered the possibility that a hypothetical alertness-modulation of the contraction of scalp muscles following TMS may have contributed to the alertness-modulation of TEP amplitude. To address this hypothesis, we compared the amplitude of the ICA component of the scalp muscle, which was removed during the first stage of ICA cleaning (Rogasch et al., 2014), between θ/α-defined awake and drowsy states. No amplitude difference was observed between awake and drowsy trials (see Fig S4), ruling out the possibility that TEP cortical-reactivity changes between awake and drowsy states were due to a change in the intensity of scalp muscle contraction.

### Statistical analysis

Paired samples t tests were used to compare behavioural and neural summary measures between θ/α-defined awake and drowsy states. Pooled variance was used to calculate Cohen’s d, with 0.2 indicating a small effect size, 0.5 a medium effect size, and 0.8 a large effect size (Cohen, 1988). For a similar comparison of summary measures across Alertness Levels 1-5, a one way repeated measures ANOVA was carried out with linear as well as non-linear contrasts. Huynh-Feldt correction was used when Mauchly’s test indicated violation of the assumption of sphericity. Partial η^2^ was calculated as an effect size in ANOVA tests, with 0.01 indicating a small effect size, 0.06 a medium effect size, and 0.14 a large effect size (Cohen, 1988). Shapiro-Wilk’s test was used to assess normality of the distribution before running parametric tests. Square-root or log10 transform were used to normalize skewed data. When transformations failed, non-parametric statistical tests were used, such as Wilcoxon’s signed-ranks test instead of a paired samples t test, and the Mann-Kendall trend test instead of a one-way repeated measures ANOVA for linear contrasts across Alertness Levels 1-5. A Jonckheere-Terpstra trend test was used to assess a linear change of single trial response variance across Alertness Levels 1-5, followed up by planned contrasts using a Mann-Whitney test. For any significant main effects, Bonferroni–Holm multiplicity correction (Holm, 1979) of p values was carried out to account for multiple follow-up comparisons between baseline Alertness Level 1 and the other four levels. Non-parametric Spearman’s rank order correlation test was used to assess for an association between single-trial MEP and TEP responses, with the Bonferroni–Holm correction of p values (Holm, 1979). Aiming to avoid sample size effects when comparing pooled single-trial data across different levels of alertness, the trial number was matched by randomly selecting 936 trials for each Alertness Level, which was the minimum number observed in one of the conditions. Statistical analyses were carried out using Matlab and IBM SPSS (v22) software packages.

## Acknowledgments

The study was supported by the Wellcome Trust (WT093811MA to TAB), the EU International Research Staff Exchange Scheme – IRSES (612681 to TAB & JBM), and the Australian Research Council (ARC) Centre of Excellence for Integrative Brain Function (CE14010007). JBM was supported by an ARC Australian Laureate Fellowship (FL110100103). We thank Dr Corinne Bareham, Abbey Nydam and Nicholas Bland for their help with data acquisition.

---

**Figure S1.**
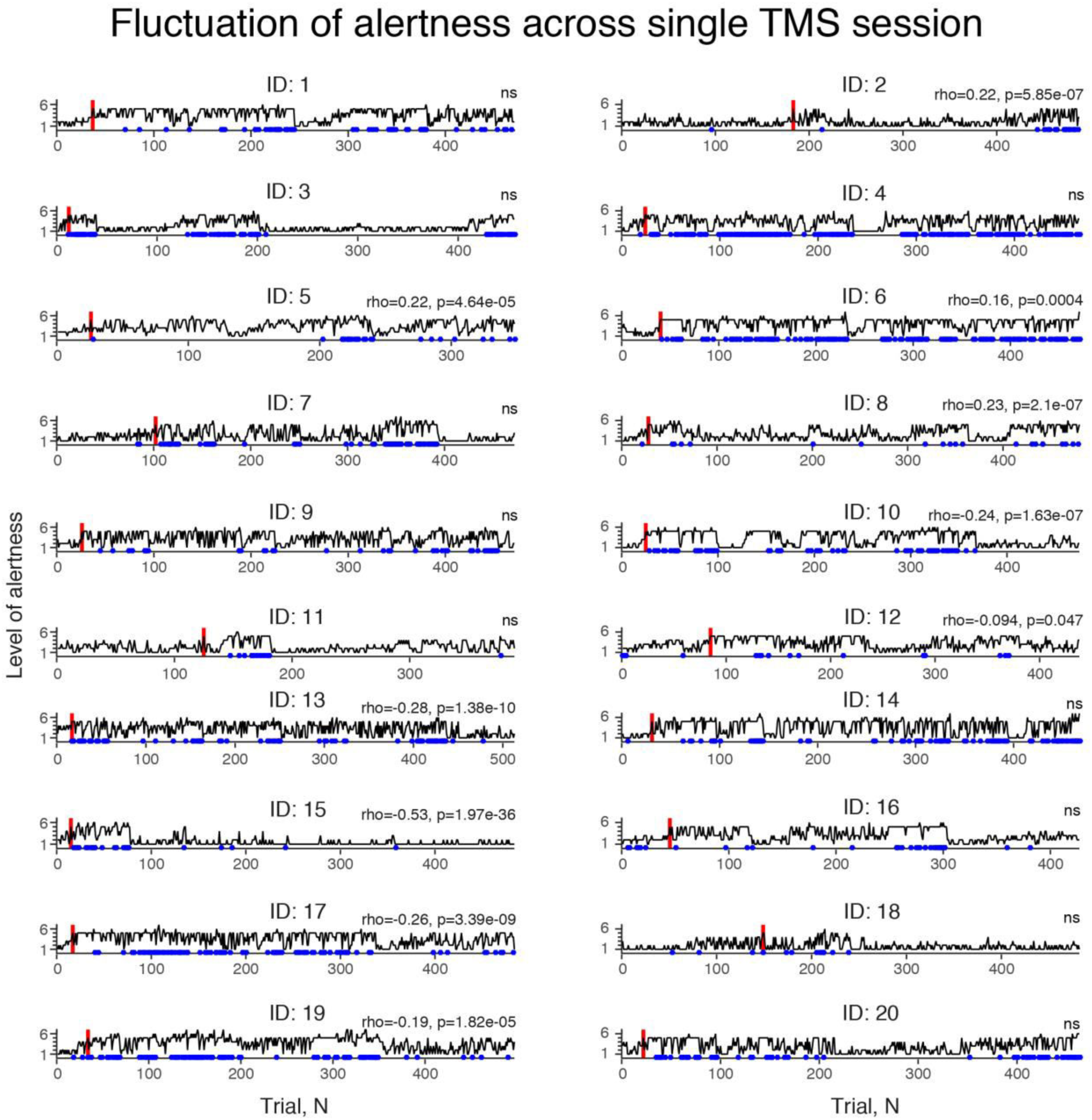
Individual differences in levels of alertness across a single TMS session. Visualisation of the alertness level (vertical axis) shown for the entire TMS testing session (horizontal axis). Each subplot represents a different participant (indicated by ID number). Red vertical lines depict the first trial within a session scored as Alertness Level 5. Blue dots indicate unresponsive trials. Results of Spearman rank order correlation tests between Alertness Level and trial number are presented in the top right corner of each subplot (ns=not significant).

**Figure S2.**
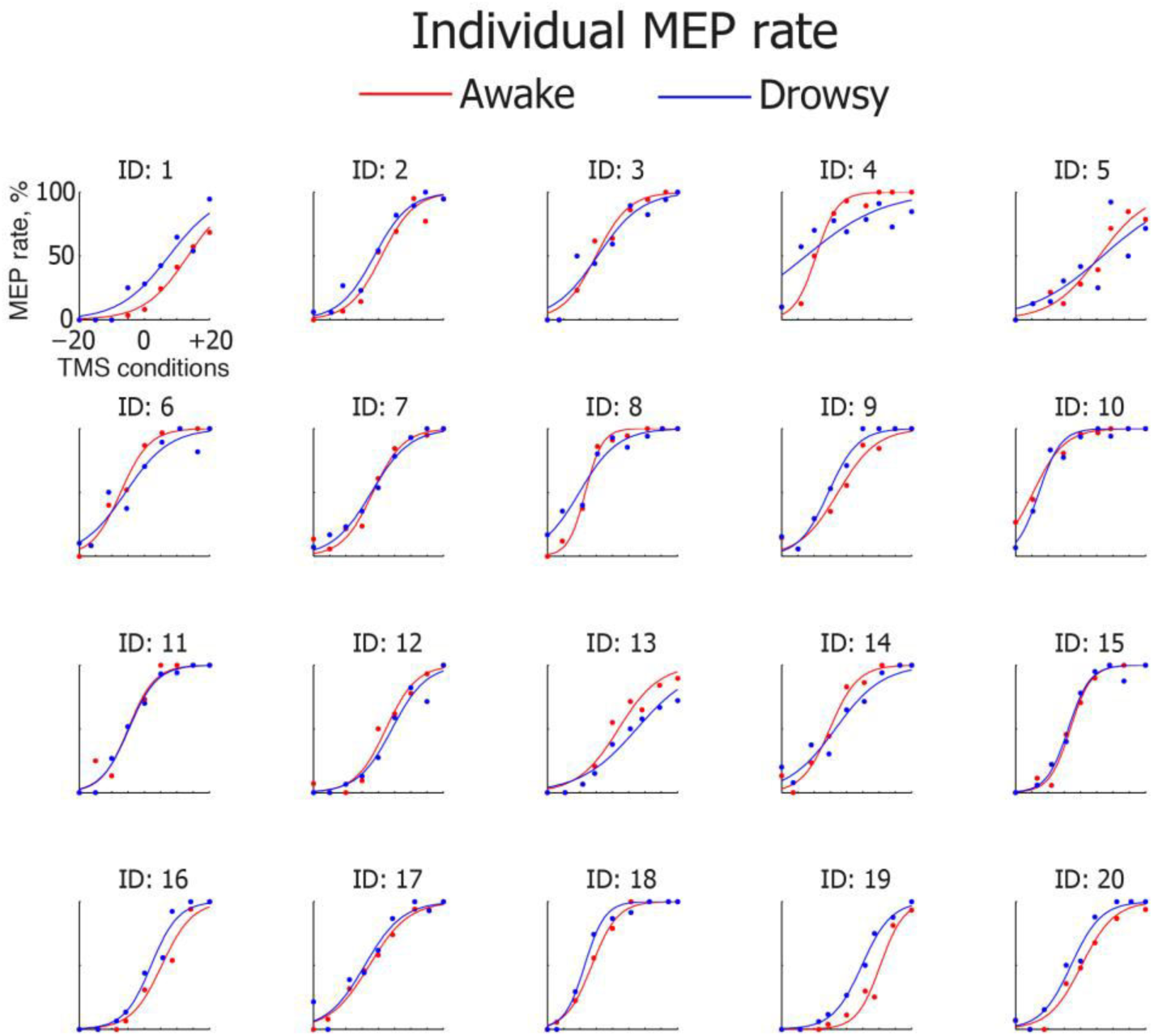
Individual rate of motor evoked potentials (MEPs) as a function of transcranial magnetic stimulation (TMS) intensity in θ/α-defined awake and drowsy states. Percentage of trials with MEPs above threshold value of 50μV, calculated separately for the awake and drowsy trials across 9 TMS conditions centred on individual motor threshold (0%). Sigmoidal functions are fitted to the awake (red) and drowsy (blue) conditions separately for each individual (N=20).

**Figure S3.**
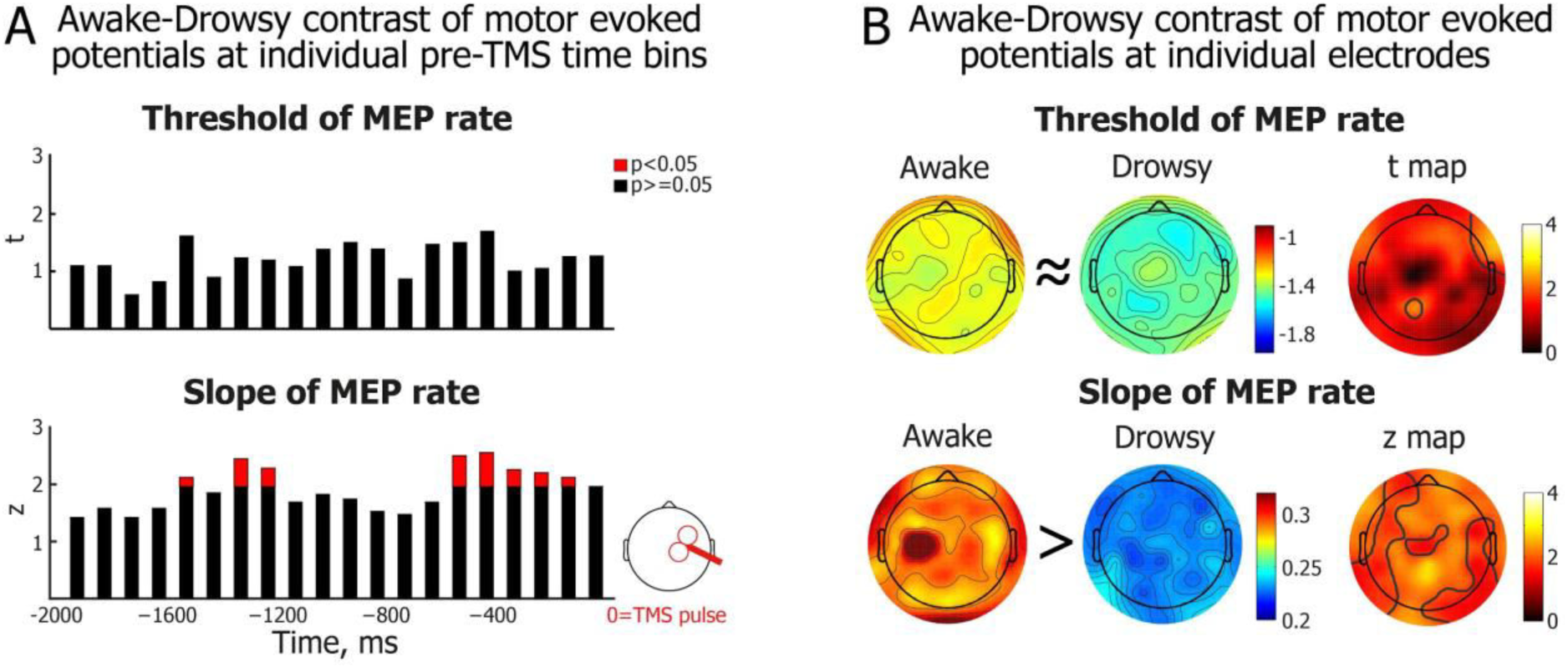
Temporal and spatial spread of the alertness-dependent modulation of motor evoked potentials. (A) Difference of the MEP sigmoid thresholds (upper panel) and slopes (lower panel) between θ/α-defined awake and drowsy conditions. Alertness states were measured and contrasted separately for each of the 20 time bins in steps of 100 ms across a 2000 to 0 ms pre-TMS time window. Electroencephalography (EEG) spectral power was averaged over all electrodes. (B) Difference in MEP sigmoid thresholds (upper row) and slopes (lower row) between θ/α-defined awake and drowsy conditions. Alertness states were measured and contrasted separately for each of the 63 EEG electrodes. EEG spectral power is averaged over a −2000 to 0 ms pre-stimulation time window.

**Figure S4.**
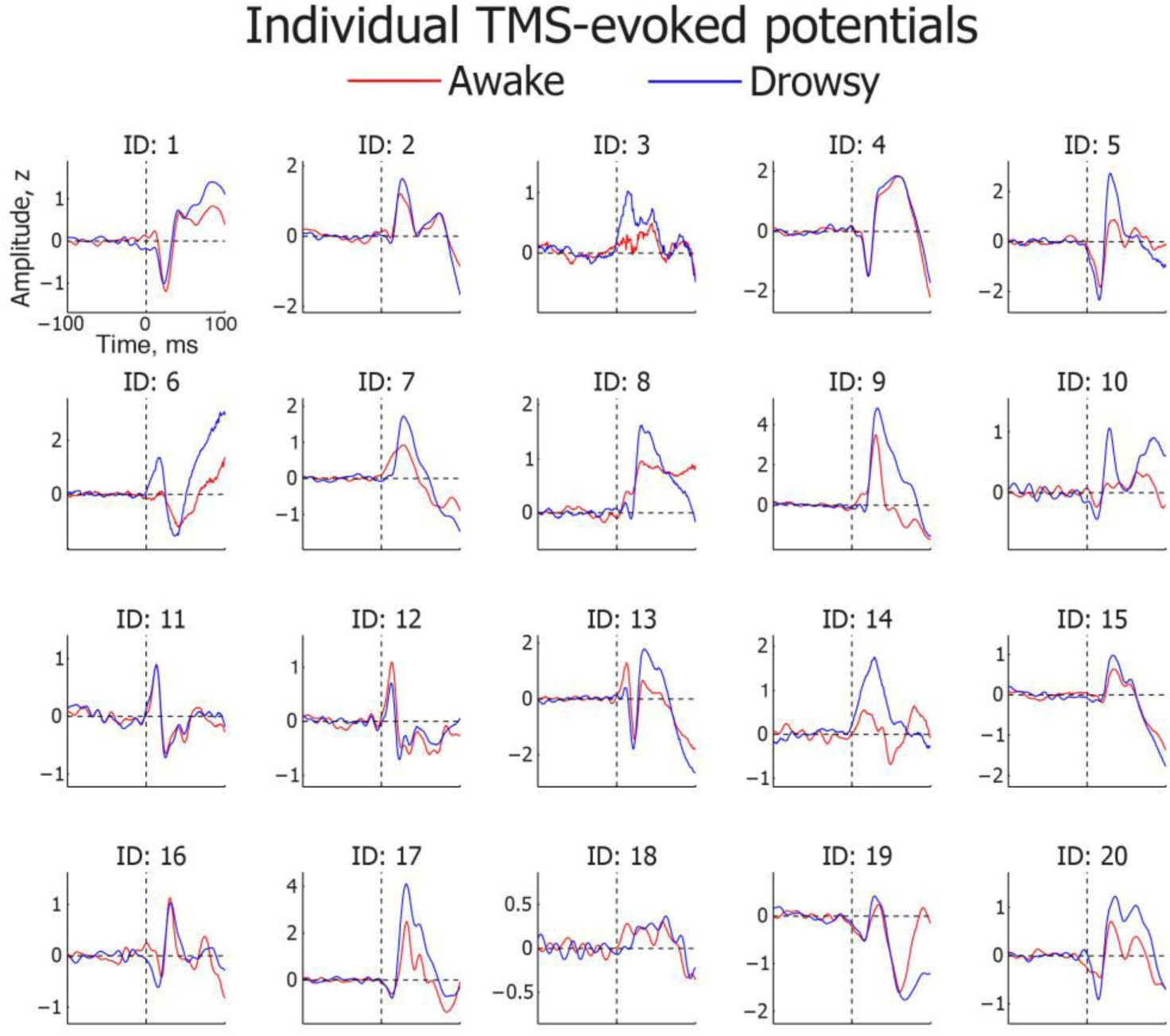
Individual transcranial magnetic stimulation-triggered cortical reactivity potentials (TEPs) in θ/α-defined awake and drowsy states. TEPs averaged across 4 EEG electrodes within a region-of-interest (ROI) beneath the TMS coil. Awake (red) and drowsy (blue) TEP waveforms are depicted separately for each individual (N=20).

**Figure S5.**
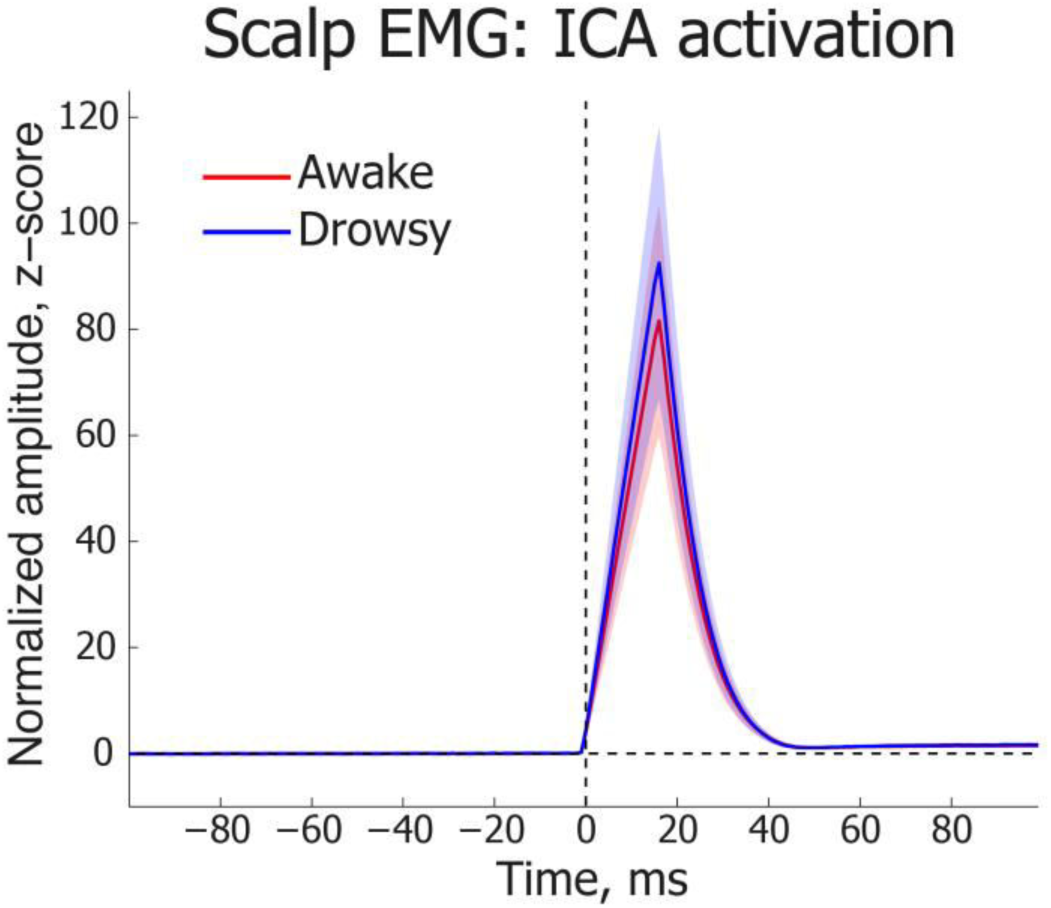
TMS-evoked scalp muscle activity in θ/α-defined awake and drowsy states. Electromyography (EMG) waveforms depicting TMS-evoked scalp muscular contraction artefact identified during independent component analysis (ICA). After identification of this component, trials were split between awake (red) and drowsy (blue) conditions. Data shown here are averaged across 16 participants in whom the artefact could be identified. Error shading depicts standard error of mean (SEM). There was no significant difference between the two conditions.

**Figure S6.**
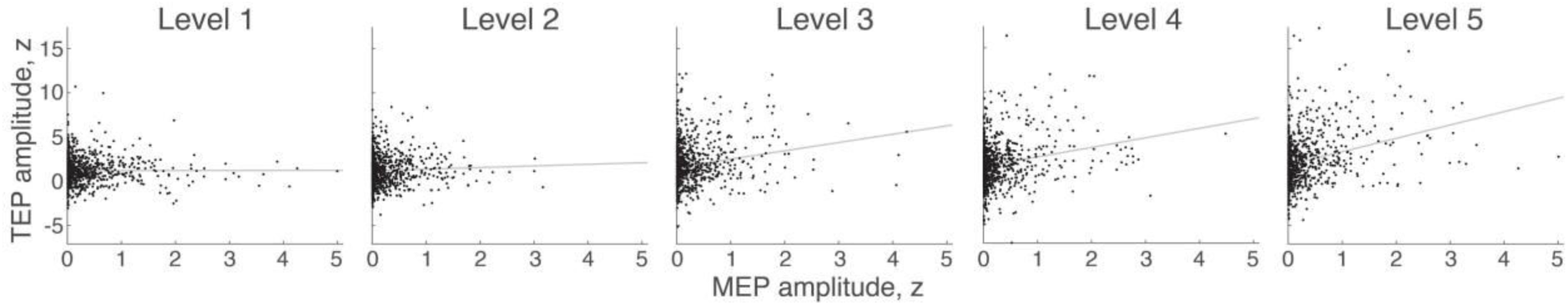
Correlations between MEP and TEP amplitudes at different levels of alertness. Individual dots in the scatter plots depict single trial amplitudes of MEP and TEP responses. Z-scored amplitude values are shown, and least-squares lines are plotted to visualize associations that were assessed using a Spearman rank order correlation test.

